# HBV pgRNA can generate a circRNA with two junction sites

**DOI:** 10.1101/2020.05.14.095273

**Authors:** Min Zhu, Zi Liang, Jun Pan, Xiaolong Hu, Xing Zhang, Renyu Xue, Guangli Cao, Chengliang Gong

## Abstract

To verify whether Hepatitis B virus (HBV) can form circRNA, the circRNA sequencing results of the HBV-positive HepG 2.2.15 hepatocarcinoma cell line were compared with the HBV genome. A novel circRNA, named as HBV_circ_1, was mapped to the 489–2985 nt region of the HBV genome (GenBank accession No. KU668446.1), which was derived from HBV pgRNA and has two junction sites. A partial fragment of the pgRNA (corresponding to HBV DNA genome 489-3,182-2,985nt) was resected and then connected to form a junction site of HBV_circ_1 (corresponding to HBV DNA genome 489/2985nt). The 5’-terminal region (corresponding to HBV DNA genome 1,820-1,932nt) of pgRNA was repeated with the region of pgRNA at the 3’-terminal region. Another junction site of HBV_circ_1 (corresponding to HBV DNA genome 1820nt) was formed in a manner similar to DNA homologous recombination mediated by repeats of pgRNA. Moreover, this RNA homologous recombination mediated by the repeats of pgRNA does not have tissue specificity and genus specificity. Reverse transcription PCR, northern blotting, and tissue in situ hybridization confirmed the existence of HBV_circ_1 in HepG2.2.15 cells and HBV related hepatocellular carcinoma (HCC) tissue. This study expands our understanding of circRNA generation mechanisms, and provide a new perspective for understanding the molecular pathogenesis of the virus.

## Introduction

Hepatitis B virus (HBV) is contagious, which can cause the body’s immune response to cause pathological damage to liver cells or tissue. While there is an approved vaccine, HBV infection remains a major global public health challenge. Approximately 2 billion patients are infected by HBV, of which 350 million are chronically infected with HBV and remain at a higher risk of developing liver cirrhosis and hepatic cellular cancer (HCC)[1, 2]. As a result, the pathogenesis of HBV has been a popular topic of study for many years. HBV is a small DNA virus that belongs to the *Hepadnaviridae* family, which is responsible for acute and chronic hepatitis in humans, which can lead to cirrhosis or liver cancer. The genomic material of HBV is circular, partially double-stranded DNA that is only 3.2 kb in size. HBV covalently closed circular DNA is transcribed by the cellular polymerase II machinery to produce viral RNAs. HBV transcription begins from different transcription start sites on the HBV genome, but it ends at a common transcription termination signal (Figure 1A). Thus, HBV transcripts differ in their 5’ termini, but share common 3’ terminal sequences. These HBV transcripts include RNAs of 2.4 and 2.1 kb that encode different forms of the hepatitis B surface antigen proteins, a 0.7-kb HBx RNA that encodes HBx protein, and two RNAs longer than the genome length (3.5-3.6 kb), known as precore RNA (pcRNA) and pgRNA[3]. It has been reported that signal transduction and the cell cycle of hepatocytes can be directly or/and indirectly regulated by HBV during the occurrence and development of HBV-related HCC. For years, the occurrence and development of HBV-related HCC have been extensively studied by transcriptomic, proteomic, metabonomics, and gene expression regulation approaches, and increased understandings has been obtained regarding the occurrence and molecular mechanism of HBV-related HCC[4–8]. Previous studies have shown that multiple paths of HBV are involved in the occurrence of HCC. It was found that HBV DNA could be integrated into the host genome, which may precede clonal tumor expansion and induces genomic instability, eventually leading to HCC[9, 10]. Prolonged expression of the viral proteins, including HBx and large envelope proteins can regulate cell death, proliferation, and signaling pathways, which are carcinogenic factors[11–13]. Moreover, HBx and HBc proteins can induce epigenetic modifications targeting the expression of tumor suppressor genes to promote the development of HCC[14, 15]. Kojima et al. showed that HBx expression interferes with telomerase activity to play a role in the carcinogenesis of HCC[16]. Previously, the pathogenesis of HBV-related HCC mainly focused on the regulation of gene expression, which encodes proteins. In recent years, it has been reported that non-coding RNAs do not encode proteins, but can regulate the expression of genes at the epigenetic, transcriptional, and post-transcriptional levels, thus participating in the pathophysiological process of HBV-related HCC[17–19].

**Figure 1.**
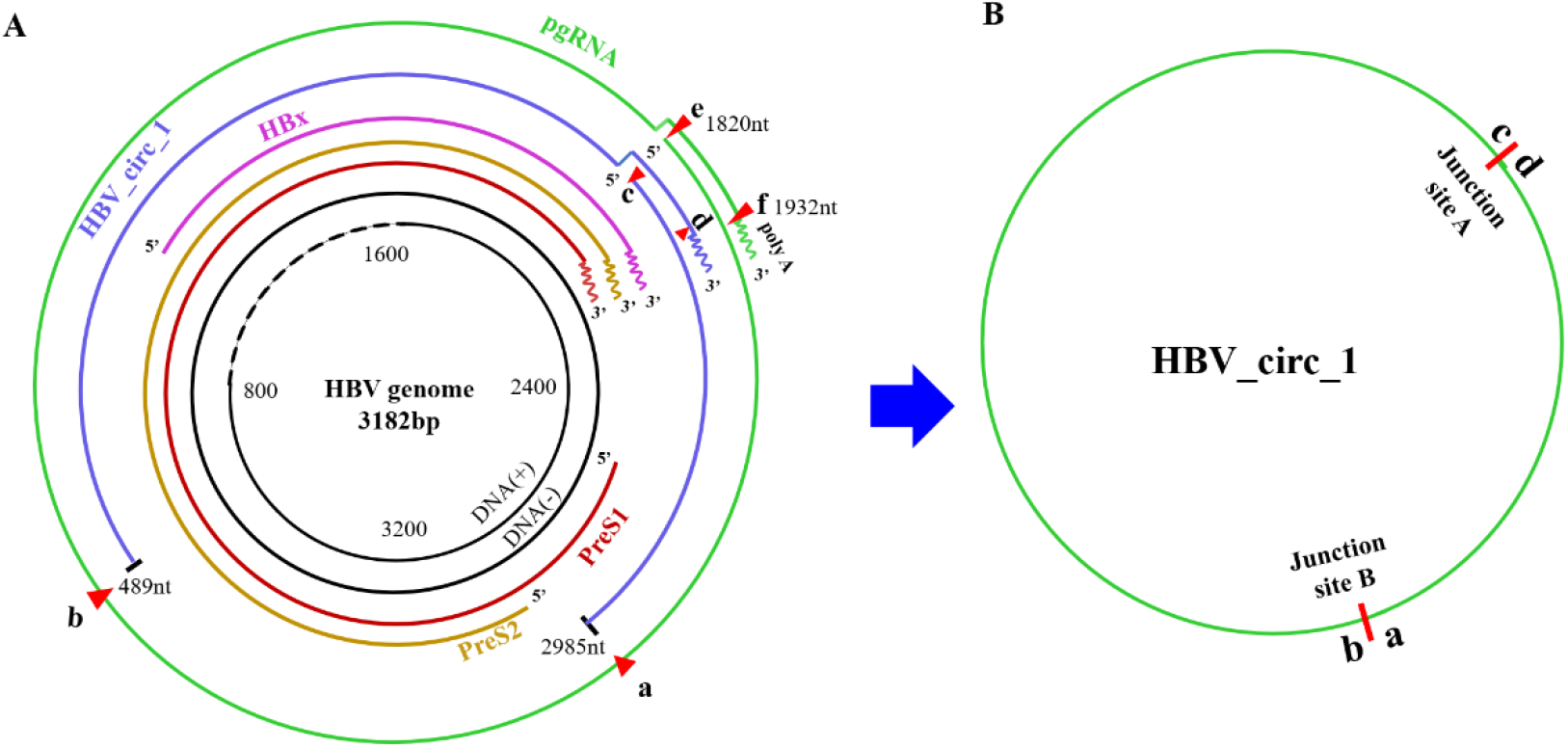
Formation mechanism of HBV_circ_1. **A**. The diagram of HBV genome and its transcripts. Green line represents the transcription intermediate of HBV pre-genomic RNA (pgRNA). Blue line represents the genomic locus of HBV_circ_1 in HBV genome. **B**. a circRNA HBV_circ_1 derived from pgRNA with two junction sites (junction site A and B).

Circular RNA (circRNA) is a type of closed circular RNA, which is produced in the process of RNA splicing, and is considered to be a new type of non-coding RNA. Recently, circRNAs have become a popular research topic in the field of RNA. In addition to cellular transcripts, it has been reported that some DNA viruses can also involve circRNAs, such as Epstein-Barr virus (EBV), Kaposi’s sarcoma herpesvirus (KSHV), rhesus macaque lymphocryptovirus, human papillomavirus(HPV)[20–23]. Ungerleider et al. used the RNase R-sequence method to identify EBV-encoded circular RNAs in latent types of EBV-infected cell models, and found that the expression of these circRNAs mainly depends on the transcriptional process of EBV and the transcription level of related sites[20], suggesting that the viral circRNAs are derived from viral mRNA. In EBV and KSHV, some of the viral circRNAs were entirely derived from exons of the viral mRNA, and some retained introns, which suggested that these viral circRNAs are formed in the same way as cellular transcripts through back-splicing[20, 21]. Whether other viruses can also form circRNAs is worth exploring. Recently, a circRNA derived from HBV pgRNA was identified[24], but the formation mechanism of circRNA encoded by HBV are still unclear. In this study, a novel circRNA derived from HBV, named as HBV_circ_1, was identified by circRNA sequencing, PCR, Sanger sequencing and northern blotting. Sequence alignment showed that HBV_circ_1 derived from HBV pgRNA, containing two junction sites (named junction site A and junction site B, respectively). Junction site A was formed by RNA homologous recombination at the end of pgRNA, junction site B by splicing of pgRNA. Moreover, the repeat sequences at pgRNA ends were found to promote circRNA formation. The results not only enrich our knowledge about the HBV genome, but also provide a new perspective for understanding the molecular pathogenesis of the virus.

## Results

### HBV_circ_1 is derived from pgRNA by splicing and recombination

We previously performed RNA sequencing with the HBV-positive cell line (HepG2.2.15) using poly(A)+-selected or RNase R-treated RNA libraries. The sequencing results showed that a circRNA termed HBV_circ_1 mapped to the 489–2,985 nucleotide (nt) region of the HBV genome (GenBank accession No. KU668446.1). Transcript analysis of HBV showed that HBV transcription began from different transcriptional start sites on the HBV genome, but it ended at a common transcription termination signal. Comparing the HBV_circ_1 with the HBV transcript suggested that HBV_circ_1 with A and B junction sites originated from pgRNA (corresponding to the 1–1,932 nt and 1,820–3,182 nt regions of HBV genome) (Figure 1A, B). It was found that the 113-base sequence of pgRNA at the 5’-terminal region was repeated with the 1,820–1,932 nt region of pgRNA at the 3’-terminal region (Figure 2A), named as RS1 and RS2, respectively. We assumed that the repeat sequence of pgRNA may drive generation of junction site A of HBV_circ_1. To confirm this assumption, RT-PCR was conducted with divergent primer1-Forward/primer1-Reverse (Figure 2A, Table 1). Sanger sequencing of amplification products showed that the 5’ end and the 3’ end of the pgRNA formed junction site A crossing 1820 nt and only a repeat sequence was retained (Figure 2B). To further confirm that the repetitive sequences drove the formation of junction site A, the sequence pgRNA-eGFP containing eGFP sequence flanked by the RS1 (1820-2131 nt) and RS2 (1621-1932 nt) was synthesized by Synbio Tech (Suzhou, China) (Supplementary 1), and subcloned to vector pcDNA3.1(+) (Invitrogen, Carlsbad, CA, USA) to generate a vector pcDNA3.1(+)-pgRNA-eGFP (Figure 2C). Then, the pcDNA3.1(+)-pgRNA-eGFP was transfected into HepG2 cells, and after the extracted RNA from the transfected cells was treated with RNase R, RT-PCR was carried out with divergent primer1-Forward/primer1-Reverse (Figure 2D). Sanger sequencing of the amplification product (Figure 2E) showed that the repeat sequence of pgRNA drove gRNA to form a junction site whose flanking sequence was consistent with that of junction site A of HBV_circ-1 (Figure 2D-F). The vector pcDNA3.1(+)-pgRNA-eGFP was also transfected into other cell lines, including ovary cells of silkworm BmN, Mouse normal liver cells AML-12, breast cancer cells MCF-7, carp epithelial tumor cells EPC, human fibroblasts HDF-a. The results showed that junction site A and one repeat unit were detected in other cell lines, except BmN cells because CMV promoter on the pcDNA3.1(+) is not active in BmN cells (Figure 3A-B). To confirm the junction site B of HBV_circ_1, after the extracted RNA from HBV-positive HepG2.2.15 cells was treated with DNaseI and RNase R to remove the DNA and linear RNA contaminations, RT-PCR was performed with primers Q-HBV_circ_1-Forward/ Q-HBV_circ_1-Reverse (Table 1). Sanger sequencing of the PCR product yielded the same spliced junction site B as high-through sequencing, further confirming that HBV transcript generated circRNA. Northern blotting confirmed the presence of HBV_circ_1 at junction site B in HBV-positive HepG2.2.15 cells (Figure 4A). Moreover, Primer2-Forward and Primer3-Reverse crossing junction site A and Primer2-Reverse and Primer3-Forward crossing junction site B were used to amplify the full length of HBV_circ_1, and the PCR product was cloned into the T vector. Sanger sequencing results yielded the same sequence of full-length HBV_circ_1 (Figure 4B).

**Figure 2.**
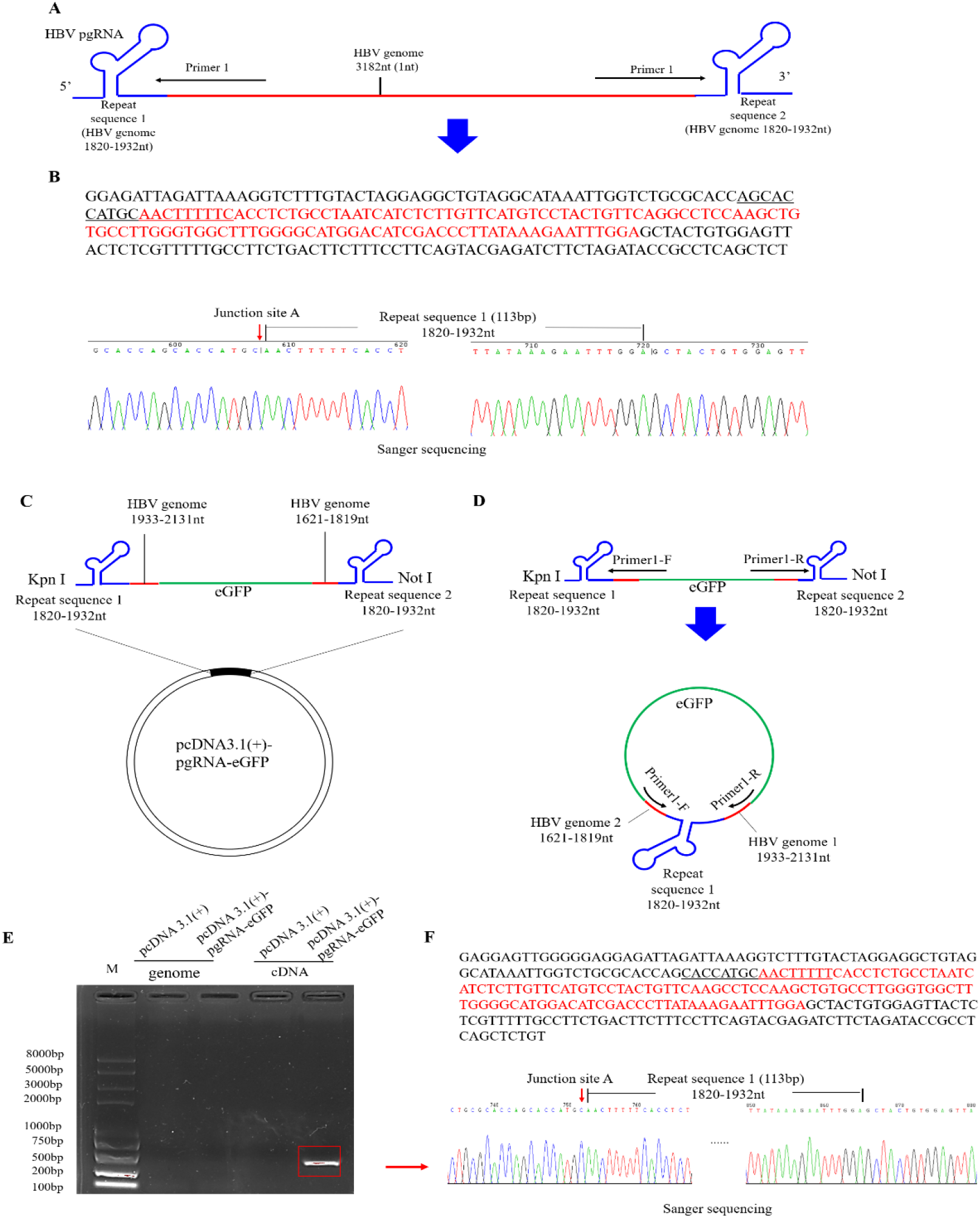
Validation of HBV_circ_1 at junction site A. **A**. The diagram of HBV pgRNA. Black arrows represent divergent primers, which are used to amplify the genome region of HBV_circ_1 containing the junction site A (HBV genome 1819-1820nt). **B**. Sanger sequencing of RT-PCR products. Upper: the repeat sequence 1 is shown as red letter. The junction site A sequence is shown as letter with underline. Lower: The peak map of Sanger sequencing. Red arrow represents the junction site A (HBV genome 1819-1820nt) by Sanger sequencing. **C**. The construction schematic diagram of the vector pcDNA3.1(+)-pgRNA-eGFP containing the 5’ and 3’ repeat sequences of pgRNA. **D**. A schematic diagram of the formation of circular RNA by the vector pcDNA3.1(+)-pgRNA-eGFP. Black arrows represent divergent primers, which are used to amplify the genome region containing the junction site A. **E**. the electrophoresis PCR products. **F**. Sanger sequencing of PCR products. Upper: the repeat sequence 1 is shown as red letter. The junction site A sequence is shown as letter with underline. Lower: The peak map of Sanger sequencing. Red arrow represents the junction site A (HBV genome 1819-1820nt) by Sanger sequencing.

**Table 1.**
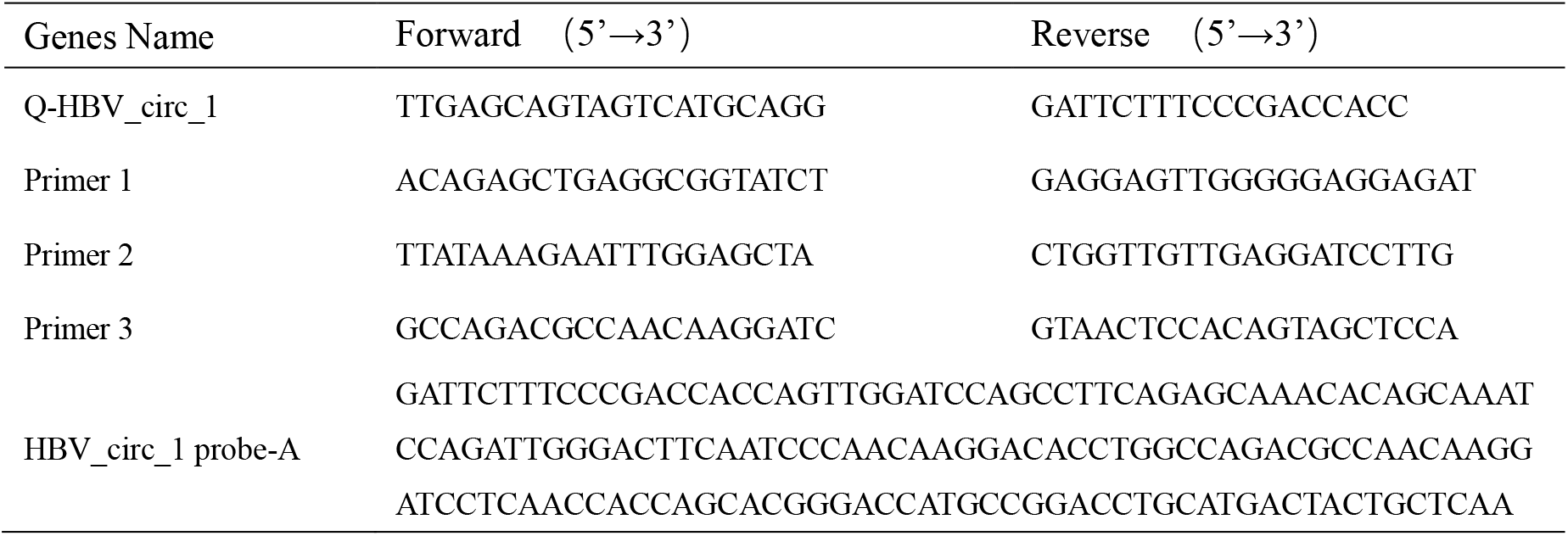
Primers and probe sequences used in this manuscript

**Figure 3.**
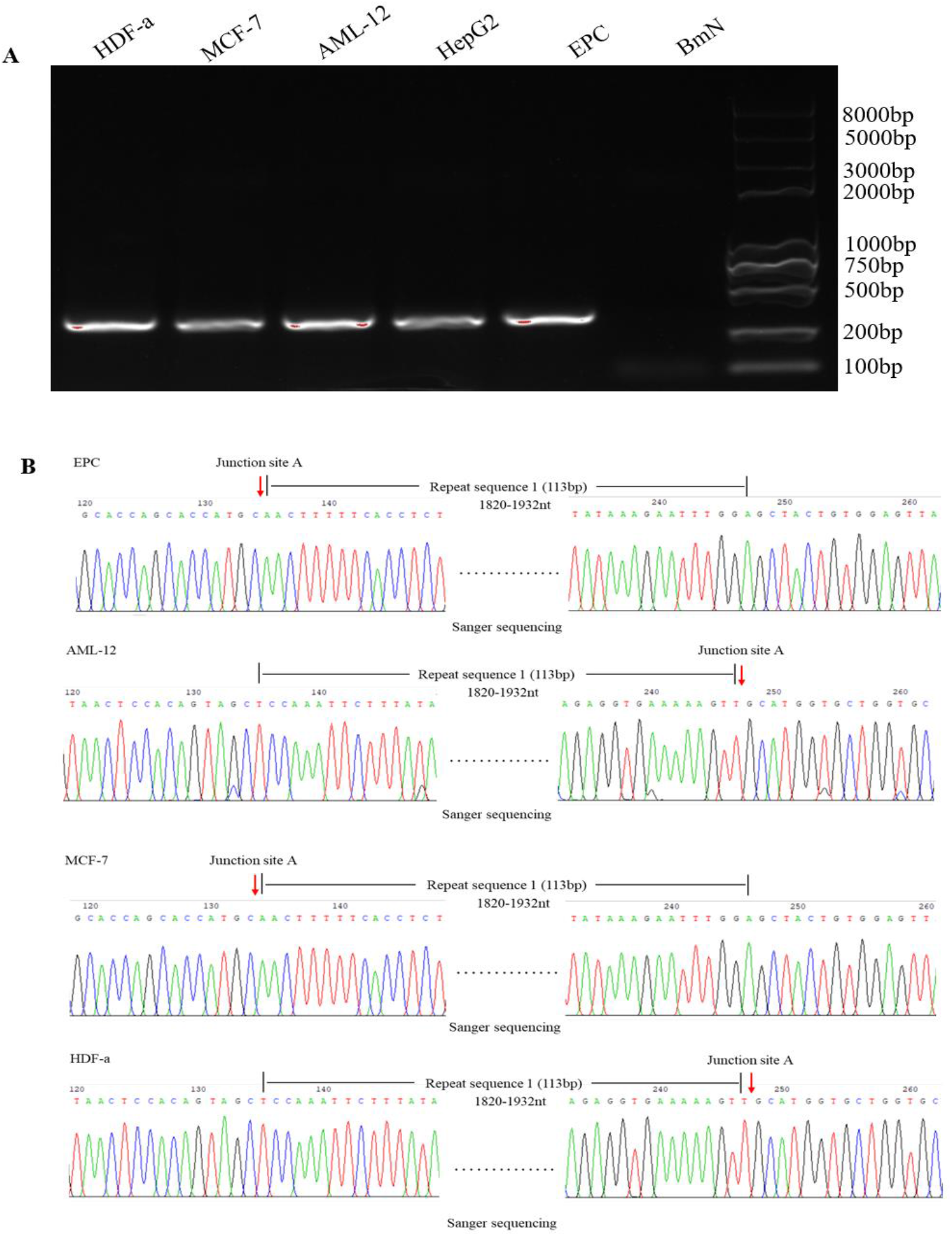
pcDNA3.1(+)-pgRNA-eGFP produced circRNA without tissue and species specificity. A, The electrophoresis of PCR products. The cells (including BmN, AML-12, Mcf-7, EPC and HDF-a) were transfected with 2μg of pcDNA3.1(+)-pgRNA-eGFP, 48 hours later, total RNA were extracted and reverse transcribed into cDNA, then PCR were performed using primer1. B-E, Sanger sequencing results of PCR products amplified from the repeative sequence unit region of generated circRNA the amplified specific bands in the detected cell lines.

**Figure 4.**
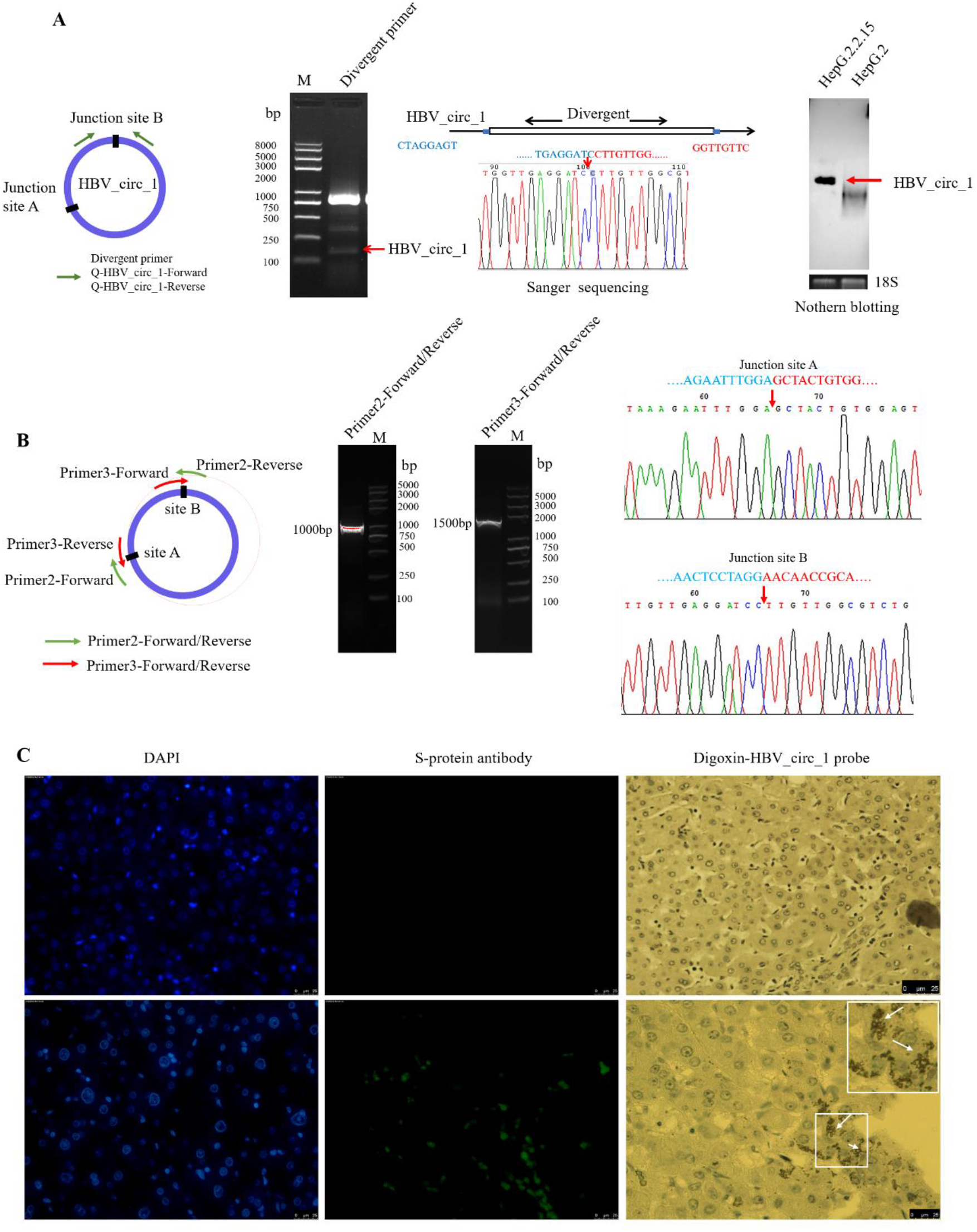
Validation of full-length HBV_circ_1. **A**, RT-PCR and Sanger sequencing validation of HBV_circ_1 at junction B. Left: the diagram of primer design for identification of HBV_circ_1. Middle: the electrophoresis RT-PCR products and Sanger sequencing of RT-PCR products. Black arrows represent divergent primers, which are used to amplify the genome region of HBV_circ_1 containing the junction site B. Red arrow represents the circRNA back-spliced junction site by Sanger sequencing. Right: Northern blotting validation of HBV_circ_1. 18S represents 18S rRNA, as the internal control. The total RNA per lane was 30 μg. DNA probes were labeled by biotin, crossing back-spliced junction site. **B**, RT-PCR and Sanger sequencing validation of HBV_circ_1. Left: the diagram of primers crossing junction site A and junction site B. Middle: the electrophoresis RT-PCR products. Right: Sanger sequencing of RT-PCR products. Red arrow represents HBV_circ_1 junction site. **C**, Validation of HBV_circ_1 in tumor tissue with hybridization in situ. Upper: HBV negative HCC tissue. Lower: HBV positive HCC tissue. Green fluorescence indicates the anti-S-protein polyclonal antibody, blue indicates DAPI, white arrow indicates the Digoxin-probe specific to HBV_circ_1 probe. The first anti-body is rabbit anti-S protein polyclonal antibody (1:1000) and the second is FITC-conjugated goat anti-rabbit IgG (1:200).

### HBV_circ_1 is present in HBV-related HCC

Our experiments confirmed that a circRNA HBV_circ_1 derived from gRNA can be found in HBV-positive cells *in vitro*. To validate the presence of HBV_circ_1 in HBV-related HCC *in vivo*, we conducted immunofluorescence and hybridization in situ experiments using tissue microarray. The results showed that no HBV_circ_1 was detected in HBV-negative tissues. In HBV-positive tissue, HBV_circ_1 was detected and mainly located in the cytoplasm, with a small amount located in the nucleus (Figure 4C). The results strongly suggested that HBV_circ_1 formed both *in vitro* and *in vivo*.

## Discussion

CircRNAs are closed circular RNA molecules, which are produced by back splicing. Increasing data have shown that a variety of organisms could generate circRNAs by back-splicing, including Drosophila[25], rice[26], silkworm[27], zebrafish[28], and humans[29]. In addition to cellular transcripts, viral transcripts were found to form circRNAs in KSHV-infected primary effusion lymphoma cells, EBV-infected cells[20, 21] or HPV16[23]. In KSHV and EBV, some of the viral circRNAs are entirely derived from exons of the viral mRNA, and some retain introns, which are formed in the same way as cellular transcripts, through back-splicing. HBV is a small DNA virus that belongs to the Hepadnaviridae family, whose genomic material of HBV is circular, partially double-stranded DNA that is only 3.2 kb in size. HBV covalently closed circular DNA is transcribed by the cellular polymerase II machinery to produce viral RNAs. HBV transcripts differ in their 5’ termini, but share common 3’ terminal sequences. These HBV transcripts include RNAs of 2.4 and 2.1 kb that encode different forms of the hepatitis B surface antigen proteins, a 0.7-kb HBx RNA that encodes HBx protein, and two RNAs longer than the genome length (3.5–3.6 kb), known as pre-core RNA (pcRNA) and pgRNA[3]. Previous studies have found that HBV pgRNA, rather than mRNA, can also form circRNA, but the way it forms has not been studied[24]. In this study, a novel HBV encoded circRNA HBV_circ_1 was found by circRNA high-throughput sequencing and preliminarily was identified by PCR, Sanger sequencing and Northern blotting. To clarifies the source of HBV_circ_1, HBV_circ_1 sequence was compared with the transcript of the HBV. The full length of HBV_circ_1 is about 2.5kb, which is longer than the HBV mRNA transcripts. Therefore, HBV_circ_1 can’t origin from HBV mRNA transcripts. It was found that HBV_circ_1 sequence can be mapped to the pgRNA, so we assume that HBV_circ_1 may originate from the pgRNA. Since HBV genes have no introns, the formation of HBV_circ_1 is different from EBV and KSHV, not by back-splicing.

Interestingly, HBV_circ_1 was found has two junction sites by sequence alignment, including junction site A and junction site B. The sequence between sites a and b on the pgRNA (corresponding to HBV DNA genome 489-3182-298nt) was resected and then connected to form a junction site B (Figure 1). Junction site B forms by a similar way as the precursor mRNA splicing, which may involve some cellular endogenous RNA endonuclease and RNA ligase. However, the specific mechanism needs further study. The pgRNA is a linear molecule whose 5’ end and 3’ end contain a repeat sequence (corresponding to HBV DNA genome 1820-1932nt), respectively (Figure 1). In this study, pgRNA 3’ and 5’ are connected to form junction site A and only one repeat sequence unit is retained. We speculate that pgRNA form junction site A mediated by pgRNA repeats in a manner similar to homologous recombination. We constructed recombinant plasmid pcDNA3.1(+)-pgRNA-eGFP with RS1-eGFP-RS2 sequences to transfected cells to confirm this speculation. It was found that the repeat sequence can promote eGFP cyclization to form junction site A. Further studies have found that as long as the promoter that controls the RS1-eGFP-RS2 sequence is active in the transfected cells, Junction site A sites retaining a repeat sequence can be detected in the pcDNA3.1(+)-pgRNA-eGFP transfected cells. This recombination pattern of the RNA we found is very similar to the classical DNA homologous recombination and does not have tissue specificity and genus specificity. Therefore, we speculate that this RNA recombination is conserved in organisms, and this phenomenon expands our understanding of circRNA generation mechanisms. In the future, we will screen proteins interacting with the repeats of pgRNA to further explore the HBV_circ_1 formation mechanism.

In summary, in this study, we showed that HBV pgRNA formed circRNA HBV_circ_1 with two junction sites, in which one junction site was formed by a similar way as the precursor mRNA splicing, and another junction site was similar to homologous recombination. Reverse transcription PCR, northern blotting, and tissue in situ hybridization confirmed the existence of HBV_circ_1 in HepG2.2.15 cells and HBV related HCC tissue. The results showed that the formation of HBV_circ_1 was different from that of cellular and viral circRNA, but the specific formation mechanism needed further study. It will be intriguing to determine whether other Hepadnaviridae viruses or small DNA viruses also encode viral RNAs. In the future, viral circRNAs may serve as important model systems for RNA biology, because they are expressed from small, well-defined operons, and these circRNA can be manipulated in the context of the viral genome. If these RNAs are found to contribute to viral tumorigenesis, they then represent a possible source for targeted therapies that may be more specific and less toxic than current therapeutic approaches.

## Materials and methods

### Cell culture and tissue

The HepG2 human liver carcinoma cell line and HBV-positive HepG2.2.15 human liver carcinoma cell line were stored in our laboratory. The SMCC-7721 human liver carcinoma cell line was kindly provided by Professor Zhou at the School of Biology & Basic Medical Science, Soochow University. The MCF-7 human breast cancer cell line MCF-7 and HDF-a human fibroblasts were kindly provided by Professor Lv at the College of Textile and Clothing Engineering of Soochow University. AML-12, a normal mouse liver cell line, was purchased from Procell Life Science & Technology (Procell, WuHan, China). All cells were maintained in DMEM/High Glucose Medium (HyClone, Logan, UT, USA) containing 10% fetal calf serum (BioInd, Kibbutz Beit Haemek, Israel) in a humidified atmosphere containing 5% CO_2_ at 37°C. The EPC carp epithelial tumor cells and BmN silkworm ovary cells was stored in our laboratory. The EPC cells were maintained in Medium 199/EBSS (HyClone, Logan, UT, USA) containing 10% fetal calf serum (BioInd, Kibbutz Beit Haemek, Israel) at 26°C, The BmN cells were maintained in TC-100 (Applichem, Gatersleben, Germany) containing 10% fetal calf serum (BioInd, Kibbutz Beit Haemek, Israel) at 26°C.

### Validation of HBV_circ_1 with Sanger-sequencing

We previously conducted circRNA sequencing of the HepG2.2.15 HBV-positive cell line using an RNase R-treated RNA library. A novel circRNA, assigned to HBV_circ_1, which mapped to the 489–2985 nt region of the HBV genome (GenBank accession No. KU668446.1), was found. Transcript analysis of HBV showed that HBV_circ_1 has two junction sites (junction sites A and B). To validate the junction sites of HBV_circ_1 with Sanger sequencing, the primers (Primer1-Forward/Primer1-Reverse, Q-HBV_circ_1-Forward/Q-HBV_circ_1-Reverse) (Table 1) were designed based on the flanking sequences of the junction sites (junction sites A and B) of HBV_circ_1 (Figure 1B). To further validate HBV_circ_1 with two junction sites, two pairs of primers (Primer2-Forward/Primer2-Reverse, Primer3-Forward/Primer3-Reverse) crossing junction sites were designed to amplify the full length of HBV_circ_1. Among the primers, Primer2-Forward and Primer3-Reverse crossed junction site A, and Primer2-Reverse and Primer3-Forward crossed the junction site B. Briefly, total RNAs were extracted using the RNAplus Kit (TaKaRa, Dalian, China) following the manufacturer’s instructions. After treatment with RNase R (Epicentre, Madison, WI, USA) to remove the linear RNAs, cDNAs were synthesised using the First Strand cDNA Synthesis kit (Transgene, Beijing, China) and a random hexamer from 1 μg of total RNA. Amplification of cDNA was conducted by PCR at 95°C for 5 min, 94°C for 50 s, followed by 30 cycles of annealing (temperature depending on the primer set used) for 50 s and extension at 72°C for 30 s, with a final extension at 72°C for 10 min. PCR products were resolved on agarose gels and the recovered PCR products were directly cloned into a T-vector for Sanger sequencing.

### Northern blotting

Detected HBV_circ_1 was further validated with northern blotting using a northern blot kit (Ambion, Austin, TX, USA) according to the manufacturer’s instructions. The biotin-labelled probe specific to HBV_circ_1 junction site B was synthesized by Sangon Biotech (Table 1). The total RNA (30 μg) was resolved using a 1% agarose-formaldehyde gel, then transferred to a Hybond-N+ nylon membrane (Roche) and hybridized with Biotin-labelled DNA probes crossing junction site B. A Biotin Chromogenic Detection kit (Thermo Scientific, Waltham, MA, USA) was used to detect the biotinylated RNAs probes.

### In situ hybridization

The HBV_circ_1 probe-A (Table 1) targeting the HBV_circ_1 B site was labelled with DIG-11-dUTP by digoxin using a Dig high primer DNA labelling kit and detection starter kit II (Roche). Hepatic tissue paraffin sections of HCC patients were treated with dewaxing and then digested with protease K. Subsequently, the coverslips were hybridized in hybridization buffer (Roche) at 37°C overnight, and the signals were detected using an Enhanced Sensitive ISH detection kit I (Boster, Wuhan, China). Cell nuclei were counterstained with 4,6-diamidino-2-phenylindole (DAPI). Finally, the images were observed using a Leica DM2000 microscope (Leica, Wetzlar, Germany).

### Immunofluorescence

HepG2 cells were seeded in 24-well plates at 1 × 10^4^ per well and cultured for 24 h. The cells were then transfected with 35 pmol biotinylated HBV_circ_1, linear HBV_circ_1 or circ_gfp. The cells were collected at 48 h post-transfection. After washing three times with 1× PBS, the cells were fixed in 4% paraformaldehyde. For immunofluorescence staining, cells were treated with the rabbit anti HBV-S protein polyclonal antibody (1:1,000), followed by CY3-conjugated anti-biotin antibodies and FITC-conjugated goat anti-rabbit IgG (1:200, Service Bio, Wuhan, China). After washing, the cells were counterstained with DAPI (1:1,000; Beyotime) and examined using a Zeiss ISM800 fluorescence microscope (Zeiss, Jena, Germany).

## Ethics approval and consent to participate

The present study was approved by the Ethics Committee of the First Affiliated Hospital of Soochow University.

## Author Approvals

All authors have seen and approved the manuscript, and that it hasn’t been accepted or published elsewhere.

