## Supplement 1 for "HBV pgRNA can generate a circRNA with two junction sites"

Supplementary data

**aactttttcacctctgcctaatcatctcttgttcatgtcctactgttcaagcctccaagctgtgccttgggtggctttggggcatggacatcgacccttataaagaatttgga**gctactgtggagttactctcgtttttgccttctgacttctttccttcagtacgagatcttctagataccgcctcagctctgtatcgggaagccttagagtctcctgagcattgttcacctcaccatactgcactcaggcaagcaattctttgctggggggaactaatgactctagctacctgggtgggtgttaatttgg GGTACCATGAGTAAAGGAGAAGAACTTTTCACTGGAGTTGTCCCAATTCTTGTTGAATTAGATGGTGATGTTAATGGGCACAAATTTTCTGTCAGTGGAGAGGGTGAAGGTGATGCAACATACGGAAAACTTACCCTTAAATTTATTTGCACTACTGGAAAACTACCTGTTCCATGGCCAACACTTGTCACTACTTTCGGTTATGGTGTTCAATGCTTTGCGAGATACCCAGATCATATGAAACAGCATGACTTTTTCAAGAGTGCCATGCCTGAAGGTTATGTACAGGAAAGAACTATATTTTTCAAAGATGACGGGAACTACAAGACACGTGCTGAAGTCAAGTTTGAAGGTGATACCCTTGTTAATAGAATCGAGTTAAAAGGTATTGATTTTAAAGAAGATGGAAACATTCTTGGACACAAATTGGAATACAACTATAACTCACACAATGTATACATCATGGCAGACAAACAAAAGAATGGAATCAAAGTTAACTTCAAAATTAGACACAACATTGAAGATGGAAGCGTTCAACTAGCAGACCATTATCAACAAAATACTCCAATTGGCGATGGCCCTGTCCTTTTACCAGACAACCATTACCTGTCCACACAATCTGCCCTTTCGAAAGATCCCAACGAAAAGAGAGACCACATGGTCCTTCTTGAGTTTGTAACAGCTGCTGGGATTACACATGGCATGGATGAACTATACAAAGCGGCCGC

cgtgaacgcccaccaaatattgcccaaggtcttacataagaggactcttggactctcagcaatgtcaacgaccgaccttgaggcatacttcaaagactgtttgtttaaagactgggaggagttgggggaggagattaggttaaaggtctttgtactaggaggctgtaggcataaattggtctgcgcaccagcaccatgc**aactttttcacctctgcctaatcatctcttgttcatgtcctactgttcaagcctccaagctgtgccttgggtggctttggggcatggacatcgacccttataaagaatttgga**

**S1. The sequence containing pgRNA repeat sequences in pcDNA3.1(+)-pgRNA-eGFP vector.** The 5’ and 3’ ends of pgRNA repeat sequences are shown as black lowercase letters, eGFP as black uppercase letters. Red lowercase letters represent the HBV genome sequence flanking the pgRNA repeat sequences.
